# An Integrative Apoptotic Reaction Model for extrinsic and intrinsic stimuli

**DOI:** 10.1101/2021.05.21.444824

**Authors:** Agustin A. Corbat, Mauro Silberberg, Hernán E. Grecco

## Abstract

Apoptosis, a form of programmed cell death central to all multicellular organisms, plays a key role during organism development and is often misregulated in cancer. Devising a single model applicable to distinct stimuli and conditions has been limited by lack of robust observables. Indeed, previous numerical models have been tailored to fit experimental datasets in restricted scenarios, failing to predict response to different stimuli. We quantified the activity of three caspases simultaneously upon intrinsic or extrinsic stimulation to assemble a comprehensive dataset. We measured and modeled the time between maximum activity of intrinsic, extrinsic and effector caspases, a robust observable of network dynamics, to create the first integrated Apoptotic Reaction Model (ARM). Observing how effector caspases reach maximum activity first irrespective of stimuli used, led us to identify and incorporate a missing feedback into a successful model for extrinsic stimulation. By simulating different recently performed experiments, we corroborated that ARM adequately describes them. This integrated model provides further insight into the indispensable feedback from effector caspase to initiator caspases.

## Introduction

Four hundred years ago, Johannes Kepler used Tycho Brahe’s unprecedented accurate measurements of celestial bodies to devise geometrical rules to describe their motion. Later, by means of Newton’s laws of motion and universal gravitation the trajectories of celestial bodies could be predicted upon any initial condition. Nowadays, in systems biology a plethora of different species have been identified and their interactions pinpointed to describe their behaviour. Through mass action law the variations in concentrations can be quantitatively predicted once kinetic rates are adequately found and interaction between different scales are considered [1].

Modeling enables an easy way of conveying prior knowledge and hypotheses as well as predicting novel experimentally testable behaviour [2]. As they reflect our knowledge of the system, these evolve when more information is uncovered. Many reaction networks are initially simple block diagrams where only influences between species are described. When an increased level of detail regarding interactions is achieved, these models evolve into systems models allowing for recognition of repeated motiffs [3]. Many experimental [4, 5, 6] and statistical tools have been implemented to detect interactions between species [7, 8]. Kinetic rates quantifying these interactions are introduced generating numerical models [9].

Caveats of model design include the difficulty of intuitively interpreting those consisting of many interconnected motifs and the vastness of parameter space [10]. Modularity, when possible, simplifies interpretability and experimental design by isolating functionality and determining which species need to be measured or perturbed [11, 12, 13]. Although efforts have been allocated to adequately find model parameters [14], some works point towards the importance of network topology [15, 16]. Additionally, large models are usually hard to extrapolate to different experimental conditions and thus lack acceptance.

Ideally, aggregating data of previous and new experiments can be used to iteratively evaluate model predictions and refine their topology as well as constrain their parameters [17, 18, 19]. However, in many cases, models are designed de novo to describe results of new experiments or qualitative predictions of expected behaviour without taking into consideration whether previous verified predictions still apply [20]. This has been made evident while studying the apoptotic signaling cascade. While the reaction network that composes this cascade can be divided into different modules that have been studied separately [21, 22, 23] and together [18, 24, 25], the overall response results from the intertwined communication between its composing pathways.

The apoptotic cascade is well conserved from an evolutionary standpoint [26]. In mammals, apoptosis can be initiated either extrinsically through death receptors or intrinsically by generating cell stress. Although extrinsic and intrinsic pathways have many intermediaries and inhibitors, both result in the activation of a specific caspase, −8 and −9, respectively. Both converge in the effector pathway, which also leads to the activation of caspases-3 and −7, which share recognition sites (see **Figure 1A**, *Inoue et al., 2009* [27]). In our previous work we addressed the apoptotic network response to extrinsic stimuli in a theoretical and experimental manner. Briefly, we introduced the implementation of homoFRET biosensors capable of measuring integral activity of caspases (**Figure 1B**). The narrow spectral bandwidth of each biosensor made it possible for the first time to observe activity of three different caspases in each single cell [18]. By timing the moment of maximum activity of caspases belonging to extrinsic, intrinsic and effector pathways we were able to refine an existing ODE based model for the apoptotic signalling cascade without losing previous predictions. Then, we combined these biosensors into CASPAM, a triple modality reporter with near equimolar co-expression that enabled larger experimental throughput and reduced spurious biological noise in network signal delays [28].

**Figure 1:**
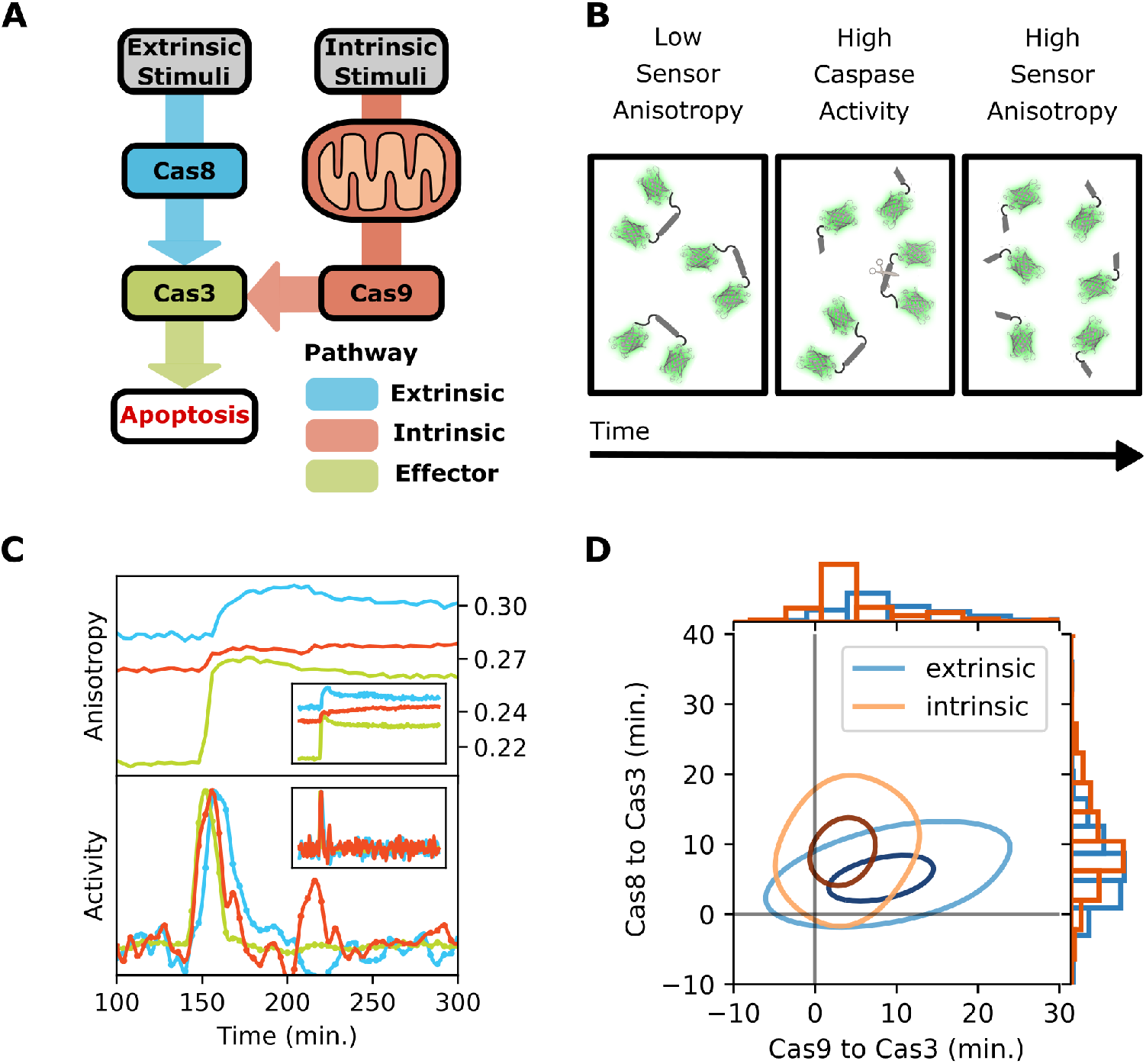
CASPAM probes timing between caspases of different pathways simultaneously at single cell level by means of anisotropy fluorescence. **A**. Sketch model of the apoptotic signalling cascade showing how information flows from stimuli, through the different initiator pathways to effector pathway and culminating in apoptosis. **B**. Biosensors are synthesized as dimers by the cell and then cleaved by a specific caspase once it is active. When in dimeric state, fluorophores excited by polarized light can transfer their energy through FRET and emit depolarized light yielding low anisotropy. When biosensors are cleaved, FRET is interrupted and emission is more polarized resulting in higher anisotropy. During cleavage, anisotropy signal is shifting and caspase activity is measured. **C**. Anisotropy curves corresponding to a cell undergoing apoptosis (upper) and the inset shows the complete curve from the 15 hr. experiment. Each color corresponds to a different biosensor for a different caspase (extrinsic-blue, intrinsic-red and effector-green). Below, the calculated normalized activity for each of the anisotropy curves and the complete curves in the inset. **D**. Bidimensional kernel density estimation plot of time difference in maximum activity between intrinsic caspase-9 and effector caspase-3 (x-axis) and extrinsic caspase-8 and effector caspase-3 (y-axis) for extrinsic (blue; N=152) and intrinsic (orange; N=183) stimuli. Being in the first quadrant translates into the effector caspase-3 reaching maximum activity first. Each curve of each distribution envelops 34% and 68% of the data points.

In this work we harnessed the power of these biosensors to generate a cohesive dataset with intrinsic and extrinsic stimuli. We aggregated these experiments to devise a single model capable of describing both. Just like the prediction of Neptune was triggered by inconsistencies between the modeled orbit of Uranus and its motion, the incompatibility between previous model predictions and our experimental results led to identifying missing feedbacks and developing a comprehensive Apoptotic Reaction Model (ARM 1.0, following previous nomenclature, *Albeck et al., 2008* [25]). We demonstrated that this new model retains prediction capability by challenging it with preceding experiments.

## Results

### Experiments show effector caspases reach maximum activity first irrespective of the kind of stimuli

Having already quantified the dynamics of the apoptotic network upon extrinsic stimuli [18], we sought to obtain a comprehensive picture by measuring the intrinsic response. We seeded HeLa cells in LabTek wells and transfected them with CASPAM, a construct encoding three biosensors specific for different caspases [28]. Then, these cells were treated with staurosporine and imaged for 15 hours. A custom made Python pipeline was implemented in order to segment cells, track them and estimate the anisotropy value for each cell across the whole experiment (see **Materials and Methods**). Anisotropy curves were further processed into activity profiles for each caspase and the time difference of maximum activity, a quantitative observable robust to intercellular variation, was recovered (see **Materials and Methods**, **Figure 1C**). We plotted the 2D kernel density estimation showing the differences between times of each caspase (**Figure 1D**). Note that although differences are in the order of minutes, onset of apoptosis across cells is highly variable and spans over several hours (**Supp. Figure 1**).

For intrinsically stimulated cells, we observe that effector caspase is the first to reach maximum activity, as it also occurs upon extrinsic stimulation [18]. Intrinsic caspase-9 reaches its peak activity 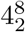 minutes later (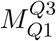, median and interquartile range) followed by extrinsic caspase-8, 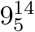 minutes later (**Figure 1D**). Correlation between time differences are the combination of two different effects. On the one hand, blue and red pairs have less dynamic range and numerical derivatives are harder to estimate resulting in higher experimental noise. On the other hand, random initial biosensor concentration, as well as other species, will cause small delays in the network which affect timing in signal propagation throughout the network.

### Current models predictions are incompatible with timing between caspases indicating missing reactions

Even though our previous model [18] was built against extrinsic stimuli experiments, it does contain the intrinsic pathway and could potentially predict experiments with such stimuli. For this purpose, we added a particular species that represents intrinsic stress, in a similar fashion as *Zhang et al., 2009* [21]. This species truncates Bid, which then interacts with other BH3 proteins initiating the signalling cascade inside mitochondria and ultimately leading to its membrane permeabilization. Due to snap action behaviour, other implementations led to identical results. Predicted activity profiles are similar to what is seen in experiments (**Figure 2B**). However, intrinsic caspase-9 is the first one to reach maximum activity in contrast to what is observed (**Figure 2B**).

**Figure 2:**
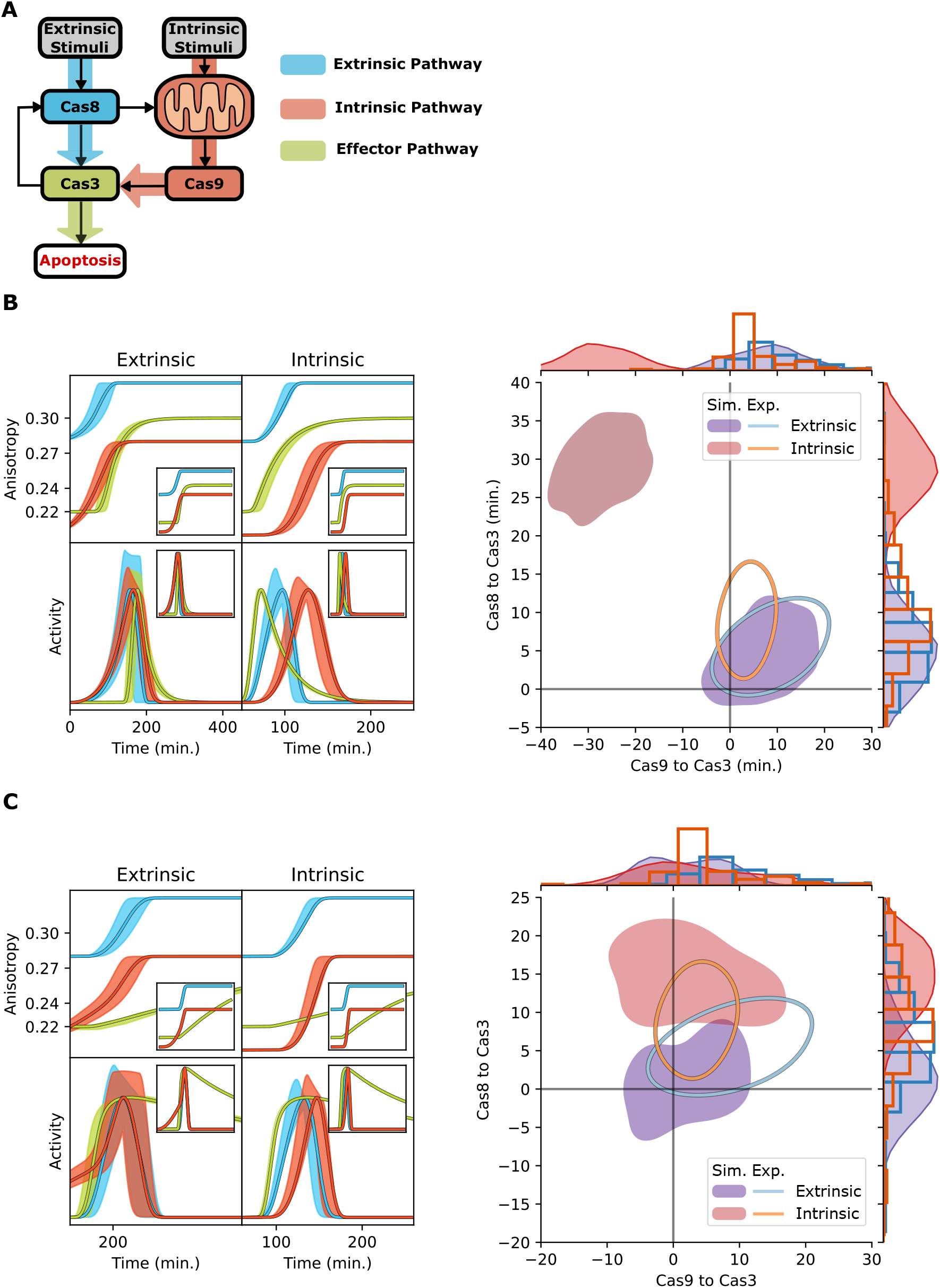
Previous models can not predict results of aggregated experiments with a single set of parameters. **A**. Sketch of the apoptotic signalling cascade with arrows detailing the reactions included in the model. Note that extrinsic caspase-8 can initiate the intrinsic pathway through mitochondria and there is a feedback between effector caspase-3 and extrinsic caspase-8 through caspase-6. **B-C**. Simulations of experimental anisotropy curves with extrinsic and intrinsic stimuli and their corresponding activity curves (left) using our previously published model (**B**, *Corbat et., 2018* [18]) and modifying parameters of a variant of the model introduced by Albeck and collaborators (**C**, Albeck et al., 2008 [25]). Filled regions describe regions where curves could fall due to variability in biosensor concentration, while the highlighted curves result from simulating the model with mid-level initial concentration of biosensors. On the right, bidimensional KDE plots enveloping 34% of the experimental data points (curves) and 66% of simulated cases (filled). **B**. Our previous model adequately predicts activity profiles as well as time differences between caspases of extrinsically stimulated cells but fails to predict order of caspases in intrinsically stimulated cells. **C**. One of the variations of the model proposed by Albeck and collaborators, is able to generate close to correct predictions of timing between caspases but fails to predict adequate activity profiles, as can be appreciated by an incomplete cleavage of effector caspase biosensor.

To understand the source of this misprediction, we began by analyzing which parameters affected the order of maximum activity between caspase-9 and −3/7 and swept them in physiological range. Specifically, species involved in reactions connecting mitochondria with those caspases, such as Cytochrome C, SMAC, XIAP, Apaf and the apoptosome. Caspase-6 was also taken into consideration as it had a possible role in controlling timing between caspase-8 and −3/7. For each sweep, we generated maps showing how timing between caspases shifted due to parameter modifications. From this analysis, we found no physiological combination of parameters that reproduced at the same time what we observed experimentally when using either kind of stimuli (**Supp. Figure 2**).

In pursuance of a model that could adequately describe all aggregated experiments, we evaluated in the same manner other available apoptotic models [24, 29, 25, 30, 31, 32]. Most of these explore different possibilities for the behaviour of species inside mitochondria. Although the region of experimental observables was accessible by a few of these models, we were unable to find a suitable combination of parameters. Timing differences were close to experimental values, when cooperativity between mitochondrial species was considered in one of the EARM variants [25]. However, in this case intrinsic caspase biosensor is never fully cleaved (**Figure 2C**). Most of these models predict that intrinsic caspase-9 is the first to achieve maximum activity, while we found that effector caspases-3/7 are actually first independently of the kind of stimuli used. Therefore, we concluded that these models were lacking a structural component, probably connecting effector and intrinsic caspases.

### Adding missing feedback reactions between effector caspases and apoptosome leads to accurate predictions

Detailed inspection of these models pointed towards a missing reaction. As previously discussed, modifying parameters in physiological ranges usually leads to small modifications [15] while topological changes might generate new behaviour. As a matter of fact, previous works and models show a positive feedback loop between caspase-9 and caspase-3/7 [33, 27, 21, 23]. Still, these models lack descriptions, and therefore predictions, of the extrinsic cascade.

By including the aforementioned feedback loop to our previous model, the order of maximum activity between caspases matches what is experimentally observed upon either stimuli. Then we refined it by tuning initial concentrations to fit experimental observations (**Supp. Model**). Unlike previous topologies in which this feedback is absent, our new model hereafter named ARM, is capable of predicting timing of maximum activity between caspases while maintaining adequate activity profiles (**Figure 3**).

**Figure 3:**
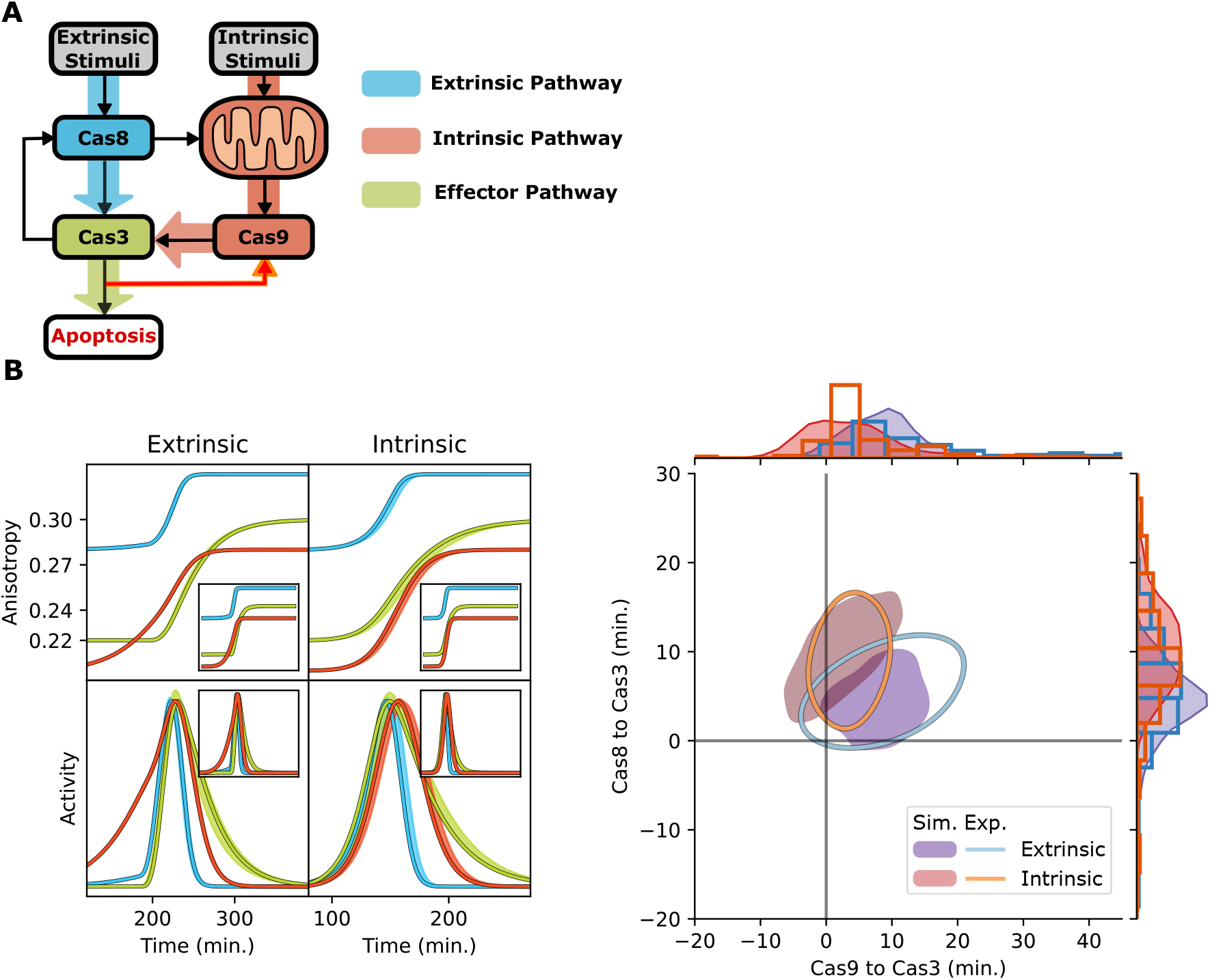
Adding missing feedback reactions to the model enables to find a set of parameters capable of predicting simultaneously experimental results when using any stimuli. **A**. Sketch of the apoptotic signalling cascade with arrows detailing the reactions included in the model. The reactions added to the model are consistent with the missing feedback between effector caspase-3 and intrinsic caspase-9 and are highlighted with a red arrow. **B**. On the left, it can be appreciated that activity profiles agree with what can be found experimentally when using any stimuli. Regarding time differences between caspases, we can see on the right that extrinsically and intrinsically stimulated and simulated cells (filled) are in close accordance with what is found experimentally (curves).

In our topological modification, cytoplasmic Cytochrome C binds to Apaf and subsequently activates it. This in turn binds and cleaves procaspase-3, instead of going through a one step conversion into the apoptosome. Once caspase-3 is active, it can bind to active Apaf and cleave its subunit, generating an active Apoptosome. The Apoptosome has a higher forward rate than active Apaf and is capable of activating more caspase-3. Moreover, these latter reactions result in a positive feedback between effector caspases-3/7 and −9 (**Figure 3A**), key for modelling the cascade dynamics as it shows the importance of effector caspases in controlling onset of apoptosis. We must highlight that the snap-action behaviour is still conserved in this model, even with lower initial concentration of apoptosome components (**Supp. Figure 3**). This is noteworthy as these species had the highest concentration.

Furthermore, analyzing the cascade as a network where activation flows from one node to another yields effector caspase-3/7 as one of the last species to be activated. Yet, experiments show that effector caspase-3/7 are the first to reach maximum activity. A possible interpretation is that initiator caspases are kept under control, while once the positive feedback loop between caspase-3/7 and −9 is reached, caspase-3/7 quickly increases its activity and removes the barriers to the other caspases.

### New Apoptotic Reaction Model preserves predictions of preceding experiments

After performing topological modifications and finding the set of parameters, we then proceeded to verify that this model still predicts the behaviour of the apoptotic cascade in well known experiments. To be specific, we expect to recover a specific relationship between onset of apoptosis with stimuli intensity and concentration of some species within the network; including interruption of signal propagation when different caspases are knocked-out.

Delay in onset of apoptosis when extrinsically and intrinsically stimulated has an inverse dependence on stimuli intensity [25,21]. This observation has been recapitulated by our model (**Figure 4A**). Additionally, the time difference between caspases is robust to modifying stimuli intensity across three orders of magnitude for any kind of stimulation (**Figure 4B**).

**Figure 4:**
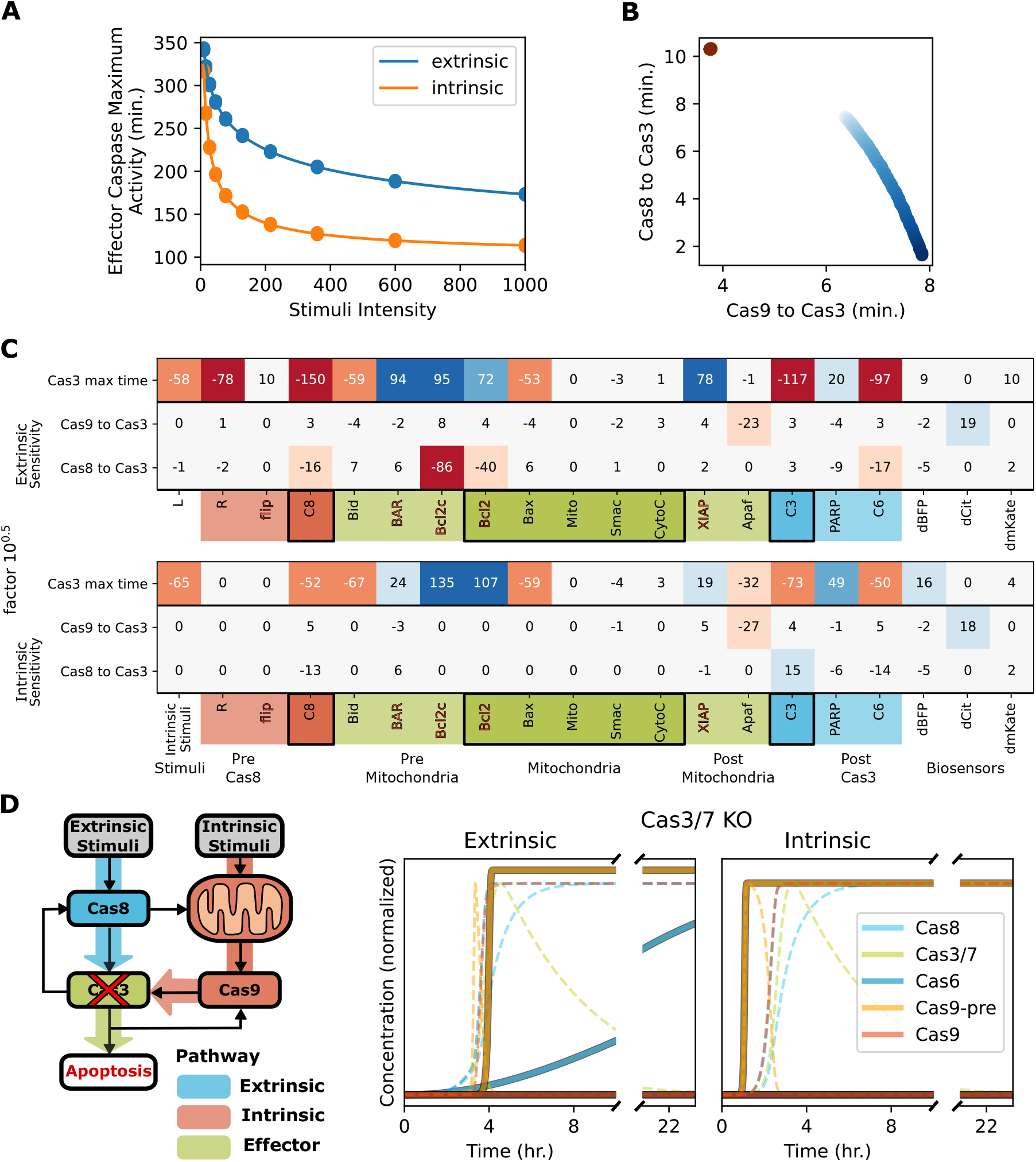
Apoptosis Reaction Model also predicts qualitative results of aggregated preceding experiments. **A.** Simulated onset of apoptosis measured as time of maximum activity of effector caspase-3 against different intensities of extrinsic (blue) or intrinsic (orange) stimuli. Previous experiments and simulations [21, 25] show the same inverse relation between onset and intensity. **B.** Scatter plot showing time difference between caspases when using different extrinsic (blue) and intrinsic (orange) stimuli intensity (darker means higher). We can appreciate that there is no variability in timing between caspases when using different intrinsic stimuli intensity, while there is a very low variation in timing when increasing extrinsic stimuli. **C.** Sensitivity analysis of observables. Model was simulated after increasing each species independently half an order of magnitude and sensitivity for each observable (effector caspase-3 maximum activity time, and time difference between extrinsic or intrinsic caspase and effector caspase) was calculated with respect to the logarithmic perturbation. Color code of sensitivity depicts sensitivity of observables to perturbations in initial concentrations. **D.** On the left, a small sketch of the model showing which nodes were interrupted by “knocking-out” effector caspases-3 and −7 from the simulation. Timelapse of activated caspases are shown on the right in order to compare with what would be expected from western blots performed at different timepoints. Dashed lines correspond to unperturbed cells while solid lines correspond to knocked-out ones. Curves are normalized to the maximum value achieved in unperturbed simulated cells. As expected, no active effector caspase is present upon either stimuli. Active extrinsic caspase-8 appears slowly only when using extrinsic stimuli as there is no feedback from caspase-6 that is never cleaved. Intrinsic caspase never achieves its final form with higher forward rates as it requires effector caspase activity. This results in a negligible cleavage of intrinsic biosensor and is in close accordance with experiments. The remaining cases are presented in **Supplementary Figure 6**.

We also performed an observable sensitivity analysis of every initial concentration in pursuance of a detailed description of their effects (see **Materials and Methods**). These consisted of onset of apoptosis, time differences of maximum activities relative to caspase 3, as well as the activity profile width measured by each sensor (**Figure 4C** and **Supp. Figure 4** and **5**). The effects on onset of apoptosis upon changing initial concentrations in an extrinsically stimulated network have been thoroughly discussed [34]. These can be summarized into how overexpression of inhibitors (FLIP, BAR, Bcl2 and XIAP) delay onset of apoptosis while caspases, receptors, and proapoptotic BH3 family members hastens it. Interestingly, SMAC, Cytochrome C or Apaf have no effect on onset. These predictions agree with experiments shown elsewhere for Bcl-2, Bid [34], FLIP and Bid [35]. Analogously, many similarities are shared in sensitivity analysis when using intrinsic stimuli instead of extrinsic stimuli. Experimental perturbations of Smac, XIAP, procaspase-3/7 concentrations have been shown elsewhere and agree with model predictions [23]. To be specific, XIAP delays onset, procaspase-3/7 hastens it and Smac does not have any effect.

We also challenged our model with experiments where subsets of caspases were knocked-out (caspase-3, −6, −7, −8 and/or −9) and cells were exposed for between 10 to 24 hours to extrinsic or intrinsic stimuli. At the end, cells were lysed and western blot was used to determine the presence of cleaved caspases [33, 27]. Although these were performed on different cell lines than the ones with which our model was devised, overall behaviour of the network is expected to be similar.

Caspase knock-outs were simulated by setting their concentration to zero or, in case of effector caspases (caspase-3/7), halving it. Propagation of the signal behaved as expected in all knock-out simulations (**Supp. Figure 6**). In close accordance with experiments, knocking-out both effector caspases (−3 and −7) resulted in no effector caspase activity being observed (**Figure 4D**). Extrinsic caspase activity is only present when using extrinsic stimuli. More interestingly, intrinsic caspase activity is negligible in both cases as feedback from effector caspases is absent. This behaviour is missing in all extrinsic stimuli based models but consistent with experimental observations previously described [23].

## Discussion

In this work we propose an integrated Apoptotic Reaction Model, to our knowledge the first that recapitulates experimental results (both our own and from others) upon either extrinsic or intrinsic stimuli. We aggregated new experiments with intrinsically stimulated cells to our previous set of observations with extrinsically stimulated ones. By means of multiplexed biosensors, defining robust experimental observables and sweeping parameter space of our model we identified a missing feedback between effector and intrinsic caspases. We verified that previously described behaviour, such as response to stimuli intensity, variation in initial concentration of specific species and knockout experiments qualitatively agreed with our model predictions.

During this process we showed that effector caspases reach maximum activity first irrespective of the stimuli used. Even though our previously modified model (based on EARM [25]) was capable of predicting this behaviour for extrinsically stimulated cells, we were unable to find a single physiological set of parameters capable of predicting these results for both stimuli. Moreover, a plethora of different models, based on experiments with extrinsic stimuli and containing the intrinsic pathway were also considered but none explained all observations. As effector caspases always reach maximum activity first, we interpreted that a feedback to intrinsic caspase was necessary and found it compatible with recent findings [33]. Adding such feedback enabled our model to recapitulate the experimental results from both extrinsic and intrinsic stimuli.

We challenged our new model against well known observations and verified that predictions agreed with experimental findings. We verified that ARM was capable of qualitatively predicting results of stimuli intensity sweeps [21,25], variations in initial concentrations [34, 35, 23] and knock-out experiments of different caspases [33, 27]. Observable sensitivity analysis is useful to develop further improvements as it yields a quick picture of which parameters can be fiddled with without affecting predictions of observables. Through integration of previous findings with our new ones, we were able to achieve a single model capable of predicting results for a broader scope of experiments.

In summary, we have shown how models can be improved constructively through data aggregation [36] and by designing tests to be applied to models based on experimental results. With this aim in mind, we devised quantitative tests that our model should comply with such as adequate prediction of timing between maximum activity of caspases and their activity profiles, as well as qualitative ones like onset of apoptosis when varying stimuli intensity or initial concentrations and topology of the network through knock-out experiments. Designing these types of tests is not straightforward as observables depend on several factors, such as cell type or growth media. A comprehensive and coherent dataset as shown here in which multiplexing simultaneously nodes from different interacting pathways [37] and reporting signals in relation to one another provides an important starting point to resolve network architecture [28, 18]. Unveiling the complete topology is crucial for performing system analysis of tumor cells responsiveness to apoptotic stimuli and mechanistic modelling of pathways that help comprehend resistance mechanisms and improve therapeutic strategies [38, 39].

## Materials and Methods

### Plasmids

CASPAM plasmid was implemented as described elsewhere [28]. This plasmid consists on three biosensors separated by flexible Gly-Ser-Gly-linkers (GGGSGGG) [40] and viral 2A sequences (P2A: ATNFSLLKQAGDVEENPGP and T2A: EGRGSLLTCGDVEENPGP). In short, the sequence reads: mCitrine-DEVD-mCitrine-P2A-tagBFP-IETD-Cerulean-T2A-mCherry-LEHD-mKate.

### Cell culture

HeLa cells for intrinsic stimuli experiments were cultured following the same procedure as before [18]. These were grown in DMEM (PAN Biotech) supplemented with 10% fetal calf serum (FCS, Gibco), 100 U/ml penicillin, 100 *μ*g/ml streptomycin, 1% L-glutamine and 1% non-essential amino-acids (all PAN Biotech) at 37 °C and 5% CO_2_ in a humidified incubator. Cells were seeded in 8 well dishes (LabTekII, Nalgene) at a density of 3×10^4^ cells. Fugene 6 (Promega) was used for transfection one day before imaging following the manufacturer’s protocol. For imaging, DMEM without Phenol red (PAN Biotech) and 0% FCS was used. For all experiments, Cycloheximide (Sigma Aldrich, Germany) was added an hour before imaging reaching a concentration of 10 *μ*g/ml. Right before imaging, staurosporine (Sigma Aldrich, Germany) was added as an intrinsic stress stimuli reaching 1 *μ*M or DMSO as control.

### Image acquisition

As previously described [41, 42], anisotropy imaging was performed by means of a custom built setup. This setup is assembled on an Olympus IX81 inverted microscope (Olympus, Germany) equipped with a MT20 illumination system. A linear dichroic polarizer (Meadowlark Optics, Frederick, Colorado, US) is placed in the illumination path of the microscope, and two identical polarizers are placed in an external filter wheel at orientations parallel and perpendicular to the polarization of the excitation light. An Orca CCD camera (Hamamatsu Photonics, Japan) was used to acquire images for each polarization and each fluorescence channel. A 20X 0.7 NA air objective and the CellR software (Olympus, Germany) were used to acquire images. Temperature and CO_2_ levels were kept constant at 37 °C and 5%, respectively, using a temperature control system consisting of an objective heater and the Stable Z specimen warmer (Bioptechs Inc., Butler, PA, USA). For the duration of the 15 hour experiment, images were acquired sequentially at every selected position of each well, at most 5 minutes apart. This experiment was repeated four times on different dates. For calibration, a dilute sample of fluorescein was used.

### Image Analysis

As before [18], dilute fluorescein images were used as normalization to correct for uneven illumination as well as for G-factor correction. Registration artifacts were corrected by means of imreg dft Python package (version 2.0 [43]). Background correction was performed by means of cellment package (version 0.2) which includes a robust estimator for the background distribution. For segmentation and tracking, a custom made python script was designed using Python 3.8, numba 0.49 [44], numpy 1.18 [45], pandas 1.0 [46], scikit-image 0.16 [47], scipy 1.4 [48] and img manager (custom built). In detail, cellment was used to determine pixels corresponding to background and after morphological operations, a mask for foreground was obtained. In order to track and separate clustered cells, cellment was implemented which leverages information from the time series such as area size and superposition in subsequent images, to apply watershed algorithm where it is necessary to separate clustered cells.

Once cells are labeled and tracked throughout the time series, sum of pixel intensity corresponding to each label are estimated for each polarization and channel. Following, anisotropy is calculated for each pair of images, and anisotropy curves are built for each object found in the images and each channel, through

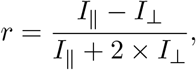

where *I*_||_ and *I*_⊥_ stand for the parallel and perpendicular measured fluorescence intensity.

By windowing these curves and fitting sigmoid functions, a coarse filtering of apoptotic curves was performed. The remaining curves are manually checked by observing all three channels of anisotropy, the change in eccentricity due to cells detaching the glass after apoptosis and watching the images from the time series. 183 apoptotic time series were obtained in the end.

### Anisotropy signal analysis

As thoroughly described in our previous work (Corbat et al., 2018), the derivative of the proportion of biosensors in monomeric state was used as a proxy for activity, as there is a linear relation between them, and is calculated through

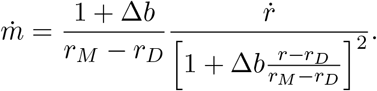

where *r_M_* and *r_D_* correspond to the anisotropy of the monomer and the dimer, respectively, and 1 + Δ*b* is the relative brightness of fluorophores in dimeric over monomeric state.

To estimate the derivative of anisotropy curves, a Savitzky-Golay filter was used, with polynomial order 2 and a window size of 20 minutes or at least 3 timepoints. As monomer and dimer anisotropy varied considerably from cell to cell, the denominator was written in terms of the normalized anisotropy curve. The maximum activity time was obtained after interpolating with splines the calculated proportional curve to monomeric fraction derivative. Seaborn 0.11 was used for kernel density estimation plots. All the code used for analysis is uploaded in a github repository named apoptoside (custom built).

### Modelling

Model implementation was done by creating a fork of the earm package developed by the Sorger Lab and is based on PySB 1.11 [24]. Our model was implemented first by instantiating Albeck’s model from EARM and modifying its initial concentrations as previously explained [18]. Subsequently, we defined the Apoptotic Reaction Model in a similar fashion, and proceeded to add the missing reactions. This model consists of 20 different species that can take different states, and 45 reactions without considering CASPAM. Our implementation, uploaded in caspase model package (Github Repository and Biomodels), takes a CASPAM parameter which, if set to True, adds 8 reactions for the 3 biosensors included. Our custom designed apoptoside package, instantiates a Model class that contains the PySB Model and generates the corresponding simulations and calculates the resulting anisotropy curves and posterior analysis resulting in activity profiles as well as timing between caspases.

When looking for a model to predict timing between caspases for extrinsic and intrinsically stimulated cells, a list of every model implemented in earm package was generated. Several parameters were swept in physiological range, such as: rates of activation of Apaf by cytochrome C, conversion of Apaf into apoptosome, inhibition of XIAP by Smac, rate of caspase-6 binding to caspase-8 and all three forward rates of caspases to their specific biosensors as well as initial concentration of caspase-6, caspase-9, Apaf and cytochrome C. For each model, every parameter was swept and timing between caspases was saved, as well as images showing the timing between caspases obtained and a sample of anisotropy and activity curves. The models closer to experimental results were manually fine tuned to obtain the best possible experimental predictions. The same procedure was applied to the model whose timing between caspases was closest to the experimental values measured. This model was obtained by using the variant from Albeck’s models that was generated in figure 11F, named albeck 11f, and increasing forward binding rate of caspase-8 to its biosensor by an order of magnitude and decreasing forward binding rate of caspase-9 to its biosensor by an order of magnitude (**Figure 2C**). In order to obtain a better adequacy between our previous model and intrinsically stimulated cells, we modified forward binding rate of caspase-8 to its biosensor by 2.5 orders of magnitude and decreasing forward binding rate of caspase-9 to its biosensor by 3.5 orders of magnitude (**Supplementary Figure 2**). For this, we sacrificed prediction timing between caspases in extrinsically stimulated cells.

Finally, once ARM was developed, we manually swept parameters in order to find the region where predictions lay closest to experimental results. After this, a grid search was performed amongst some parameters to find the best possible set of parameters. These parameters included: binding rates of all three caspases to their corresponding biosensors, Apaf to intrinsic biosensor and of caspase-3 to Apaf, and initial concentrations of biosensors, Apaf, XIAP and Cytochrome C. In order to find the best fit, timing between caspases was considered, as well as caspase activity width, defined as full width at half maximum. Target timing between caspases was of 10 minutes between caspase-9 and caspase-3 and 5.5 minutes between caspase-8 and caspase-3 for extrinsic stimuli; and 4 minutes between caspase-9 and caspase-3 and 9 minutes between caspase-8 and caspase-3 for intrinsic stimuli. Target widths were 15 minutes for caspase-3 and −8 and 30 minutes for caspase-9 irrespective of stimuli. After finding the best combination of parameters, further improvement was performed manually. Final parameters of the model can be found in **Supplementary Model** and uploaded (caspase model and Biomodels).

Stimuli sweeps were performed by varying the initial concentration of ligand (parameter: L 0) or intrinsic stress (parameter: IntrinsicStimuli) between 10 and 10^3^. For sensitivity analysis, each initial concentration was increased and decreased half an order and an order of magnitude independently, and the model was simulated and analyzed to quantify its observables (caspase-3 maximum activity time, time difference of maximum activity between caspase-8 and −3 and between caspase-9 and −3, and activity width of each caspase). Sensitivity was calculated as

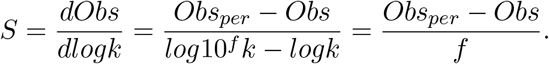

As parameter *k* was varied by an order of magnitude or half an order of magnitude, the denominator can only take the values (±0.5 or ±1).

For “knock-out” simulations, initial concentrations of corresponding procaspases (Caspase, −6, −8, −9) were set to 0. As caspase-3 and −7 are redundant in this model, results for caspase-7 are identical to caspase-3 knock-outs and this was done by halving the initial concentration of procaspase-3. Single knock out simulations were performed for caspase-3, −6, −7, −8, −9; double knock out simulations included caspase-3 and −7, −3 and −6; and triple knock-out for caspase-3, −6 and −7. For each knock-out experiment, simulation was run for 24 hours and active concentration of each caspase was saved. Active caspase concentration and substrate cleavage were assessed to determine if enough caspase is active and could possibly be detected in western blot experiments.

## Acknowledgements

We thank Dr. Sven Müller for technical assistance and Dr. Klaus Schuermann for useful discussions. We also thank Consejo Nacional de Investigaciones Científicas y Técnicas (CONICET) and University of Buenos Aires (UBA) for financial support to AAC, MS and HEG. This work was supported by the following grants: PICT 2014-3658; PICT 2013-1301; Max Planck Gesellschaft Partner Group.

## Author Contributions

Conceptualization, A.A.C. and H.E.G.; Methodology, A.A.C. and H.E.G.; Software, A.A.C., M.S. and H.E.G.; Investigation, A.A.C.; Data Curation, A.A.C. and M.S.; Writing – Original Draft, A.A.C. and H.E.G.; Writing – Review and Editing, A.A.C., M.S. and H.E.G.; Visualization, A.A.C. and H.E.G.; Supervision, H.E.G; Project Administration, H.E.G.; Funding Acquisition, H.E.G.; Resources, H.E.G.

## Declaration of Interests

The authors declare no competing interests.

## Notes

https://www.ebi.ac.uk/biomodels/MODEL2105210001

### Competing Interest Statement

The authors have declared no competing interest.

