## Supplementary Information for "An Integrative Apoptotic Reaction Model for extrinsic and intrinsic stimuli"

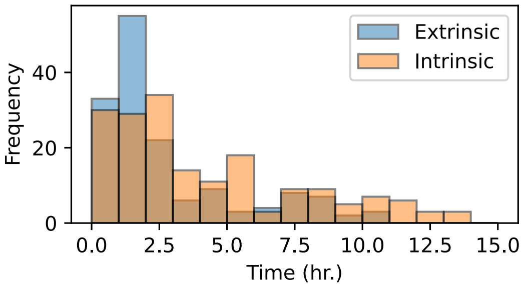


**Supplementary Figure 1: Onset of apoptosis has a lot of variability ranging from a couple hours to 15 hours.** Histogram of onset of apoptosis observed for 152 extrinsically stimulated cells (blue) and 183 intrinsically stimulated cells (orange).

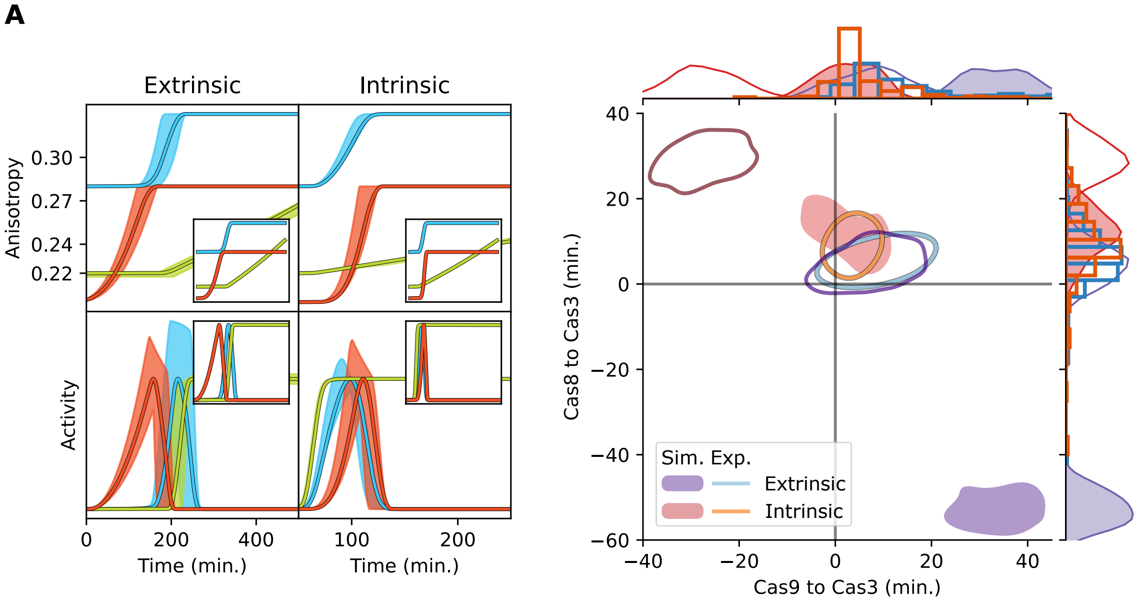


**Supplementary Figure 2: Modifying parameters of specific models improves predictions of time differences between caspases but disturbs activity profiles.** We can appreciate on the left how effector caspase activity is not enough to cleave its substrates during the 15 hours of simulation and yields abnormal activity profiles. On the right, modifying parameters can improve predictions of timing between caspases for intrinsic stimuli at the expense of not being able to predict timing for extrinsic stimuli.

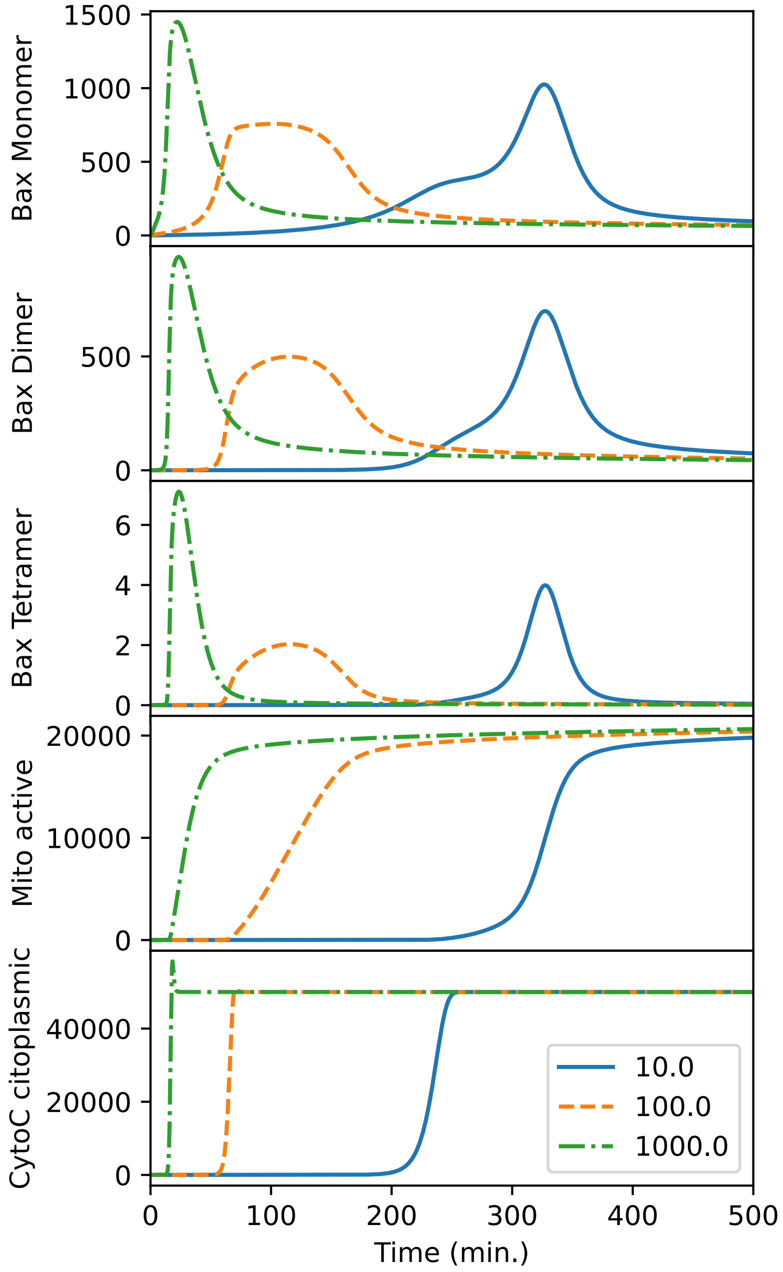


**Supplementary Figure 3: Mitochondrial contents are spilled to the cytoplasm after a few mitochondrial pores are formed.** Time lapse of concentration of different mitochondrial species after intrinsically stimulating the cells with logarithmically increasing levels of stimuli. Note that Bax dimerizes and tetramerizes at different rates, but once a few tetramers open one to three pores, cytochrome C is quickly transported to the cytoplasm. This is a key feature of the snap action behaviour.

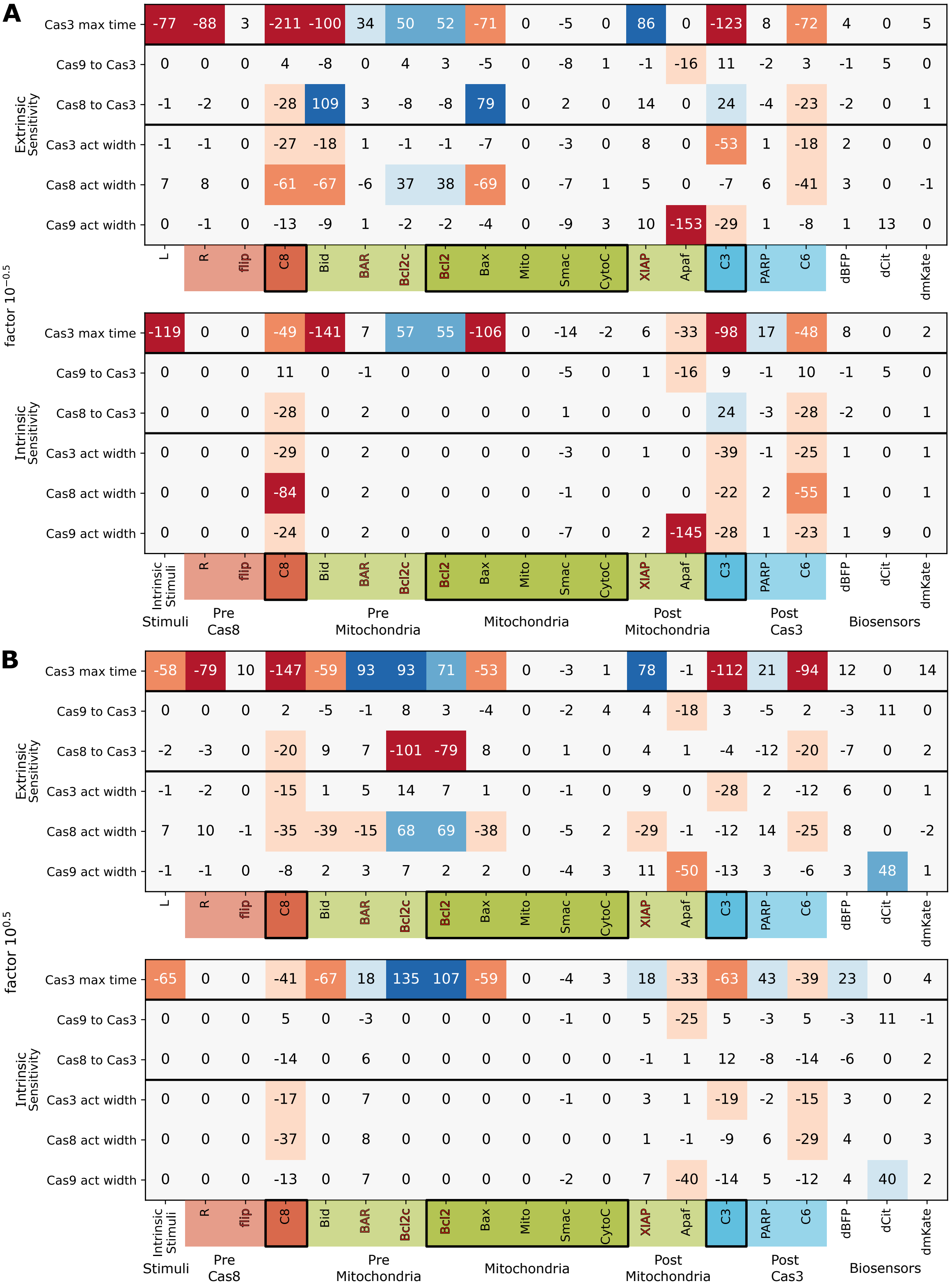


**Supplementary Figure 4: Sensitivity of observables to varying the initial concentration of each species in the model independently half an order of magnitude.** (Continued in following page)

**Supplementary Figure 4: A-B**. Sensitivity analysis of observables. Model was simulated after reducing (**A**) or increasing (**B**) each species independently half an order of magnitude and sensitivity for each observable (Effector caspase-3 maximum activity time, time difference between extrinsic or intrinsic caspase and effector caspase and caspase full width at half maximum activity) was calculated with respect to the logarithmic perturbation. Color code of sensitivity depicts which initial concentrations can be modified without producing any effect on a particular observable. With regard to intrinsic and effector caspase timing differences, these are mainly affected by Apaf. While using extrinsic stimuli, timing between extrinsic and effector caspase is reduced by increasing caspase-8 and -6, as well as, Bcl2.

**A**

factor $10^{-1.0}$

Extrinsic Sensitivity

| | L | R | flip | C8 | Bid | BAR | Bcl2c | Bcl2 | Bax | Mito | Smac | CytoC | XIAP | Apaf | C3 | PARP | C6 | dBFP | dCit | dmKate |
|---|---|---|---|---|---|---|---|---|---|---|---|---|---|---|---|---|---|---|---|---|
| Cas3 max time | -83 | -90 | 2 | -248 | 0 | 23 | 40 | 47 | -1 | 0 | -5 | 0 | 74 | 0 | -125 | 5 | -55 | 3 | 0 | 3 |
| Cas9 to Cas3 | 0 | 0 | 0 | 7 | -425 | 0 | 3 | 3 | -425 | 0 | -28 | -1 | -4 | -17 | 14 | -2 | 2 | -1 | 3 | 0 |
| Cas8 to Cas3 | -1 | -1 | 0 | -38 | 5 | 2 | -5 | -5 | 5 | 0 | 3 | 0 | 7 | 0 | 38 | -3 | -26 | -2 | 0 | 0 |
| Cas3 act width | -1 | -1 | 0 | -36 | -70 | 1 | 0 | 0 | -70 | 0 | -16 | 0 | -8 | 0 | -86 | 1 | -18 | 1 | 0 | 0 |
| Cas8 act width | 6 | 7 | 0 | -134 | -32 | -4 | 20 | 20 | -32 | 0 | -15 | 0 | 14 | 0 | -29 | 4 | -73 | 2 | 0 | 0 |
| Cas9 act width | 0 | 0 | 0 | -17 | -162 | 1 | -2 | -2 | -157 | 0 | -48 | 2 | 9 | -314 | -43 | 0 | -8 | 1 | 8 | 0 |

Intrinsic Sensitivity

| | Intrinsic Stimuli | R | flip | C8 | Bid | BAR | Bcl2c | Bcl2 | Bax | Mito | Smac | CytoC | XIAP | Apaf | C3 | PARP | C6 | dBFP | dCit | dmKate |
|---|---|---|---|---|---|---|---|---|---|---|---|---|---|---|---|---|---|---|---|---|
| Cas3 max time | -161 | 0 | 0 | -52 | -206 | 4 | 38 | 42 | -270 | 0 | -46 | -5 | 4 | -33 | -124 | 12 | -52 | 5 | 0 | 1 |
| Cas9 to Cas3 | 0 | 0 | 0 | 16 | 0 | -1 | 0 | 0 | -7 | 0 | -23 | 0 | 1 | -15 | 14 | -1 | 16 | -1 | 3 | 0 |
| Cas8 to Cas3 | 0 | 0 | 0 | -40 | 0 | 1 | 0 | 0 | 2 | 0 | 4 | 0 | 0 | 0 | 36 | -2 | -43 | -1 | 0 | 1 |
| Cas3 act width | 0 | 0 | 0 | -40 | 0 | 1 | 0 | 0 | -3 | 0 | -19 | 0 | 0 | 0 | -67 | -1 | -34 | 1 | 0 | 0 |
| Cas8 act width | 0 | 0 | 0 | -148 | 0 | 1 | 0 | 0 | -2 | 0 | -8 | 0 | 0 | 0 | -31 | 1 | -81 | 1 | 0 | 1 |
| Cas9 act width | 0 | 0 | 0 | -31 | 0 | 1 | 0 | 0 | -5 | 0 | -35 | 0 | 1 | -306 | -41 | 1 | -30 | 1 | 5 | 0 |

Stimuli · Pre Cas8 · Pre Mitochondria · Mitochondria · Post Mitochondria · Post Cas3 · Biosensors

**B**

factor $10^{1.0}$

Extrinsic Sensitivity

| | L | R | flip | C8 | Bid | BAR | Bcl2c | Bcl2 | Bax | Mito | Smac | CytoC | XIAP | Apaf | C3 | PARP | C6 | dBFP | dCit | dmKate |
|---|---|---|---|---|---|---|---|---|---|---|---|---|---|---|---|---|---|---|---|---|
| Cas3 max time | -45 | -71 | 17 | -122 | -52 | 144 | 1 | 1 | -49 | 0 | -2 | 2 | 86 | -2 | -103 | 35 | -91 | 21 | 0 | 27 |
| Cas9 to Cas3 | 0 | 1 | 0 | 2 | -4 | -3 | 350 | 425 | -3 | 0 | -1 | 4 | 65 | -17 | 5 | -8 | 2 | -5 | 17 | 0 |
| Cas8 to Cas3 | -2 | -5 | 0 | -17 | 5 | 16 | -5 | -5 | 5 | 0 | 1 | 0 | -4 | 1 | -6 | -22 | -20 | -14 | 0 | 4 |
| Cas3 act width | -1 | -2 | 0 | -10 | 0 | 12 | 70 | 70 | 0 | 0 | -1 | 0 | 61 | -1 | 8 | 4 | -8 | 11 | 0 | 2 |
| Cas8 act width | 5 | 8 | -2 | -36 | -20 | 5 | 32 | 32 | -21 | 0 | -4 | 2 | 24 | -2 | -19 | 20 | -29 | 13 | 0 | -3 |
| Cas9 act width | 0 | -1 | 0 | -6 | 2 | 6 | 187 | 67 | 2 | 0 | -3 | 2 | 300 | -35 | -9 | 7 | -5 | 6 | 112 | 1 |

Intrinsic Sensitivity

| | Intrinsic Stimuli | R | flip | C8 | Bid | BAR | Bcl2c | Bcl2 | Bax | Mito | Smac | CytoC | XIAP | Apaf | C3 | PARP | C6 | dBFP | dCit | dmKate |
|---|---|---|---|---|---|---|---|---|---|---|---|---|---|---|---|---|---|---|---|---|
| Cas3 max time | -48 | 0 | 0 | -36 | -48 | 28 | 200 | 510 | -46 | 0 | -1 | 6 | 145 | -33 | -51 | 66 | -35 | 39 | 0 | 8 |
| Cas9 to Cas3 | 0 | 0 | 0 | 4 | 0 | -7 | 0 | -2 | 0 | 0 | -1 | 0 | 61 | -23 | 4 | -6 | 4 | -5 | 15 | -1 |
| Cas8 to Cas3 | 0 | 0 | 0 | -10 | 0 | 16 | 0 | -12 | 0 | 0 | 0 | 0 | -8 | 3 | 10 | -15 | -11 | -11 | 0 | 5 |
| Cas3 act width | 0 | 0 | 0 | -13 | 0 | 15 | 0 | 21 | 0 | 0 | 0 | 0 | 48 | 1 | -15 | 0 | -11 | 7 | 0 | 4 |
| Cas8 act width | 0 | 0 | 0 | -27 | 0 | 24 | 0 | 11 | 0 | 0 | 0 | 0 | 31 | -2 | -6 | 13 | -22 | 7 | 0 | 6 |
| Cas9 act width | 0 | 0 | 0 | -10 | 0 | 14 | 0 | -24 | 0 | 0 | -1 | 0 | 291 | -32 | -10 | 11 | -9 | 8 | 105 | 3 |

Stimuli · Pre Cas8 · Pre Mitochondria · Mitochondria · Post Mitochondria · Post Cas3 · Biosensors

**Supplementary Figure 5: Sensitivity of observables to varying the initial concentration of each species in the model independently an order of magnitude.** (Continued in following page)

**Supplementary Figure 5: A-B**. Sensitivity analysis of observables. Model was simulated after reducing (**A**) or increasing (**B**) each species independently an order of magnitude and sensitivity for each observable (Effector caspase-3 maximum activity time, time difference between extrinsic or intrinsic caspase and effector caspase and caspase full width at half maximum activity) was calculated with respect to the logarithmic perturbation. Color code of sensitivity depicts which initial concentrations can be modified without producing any effect on a particular observable.

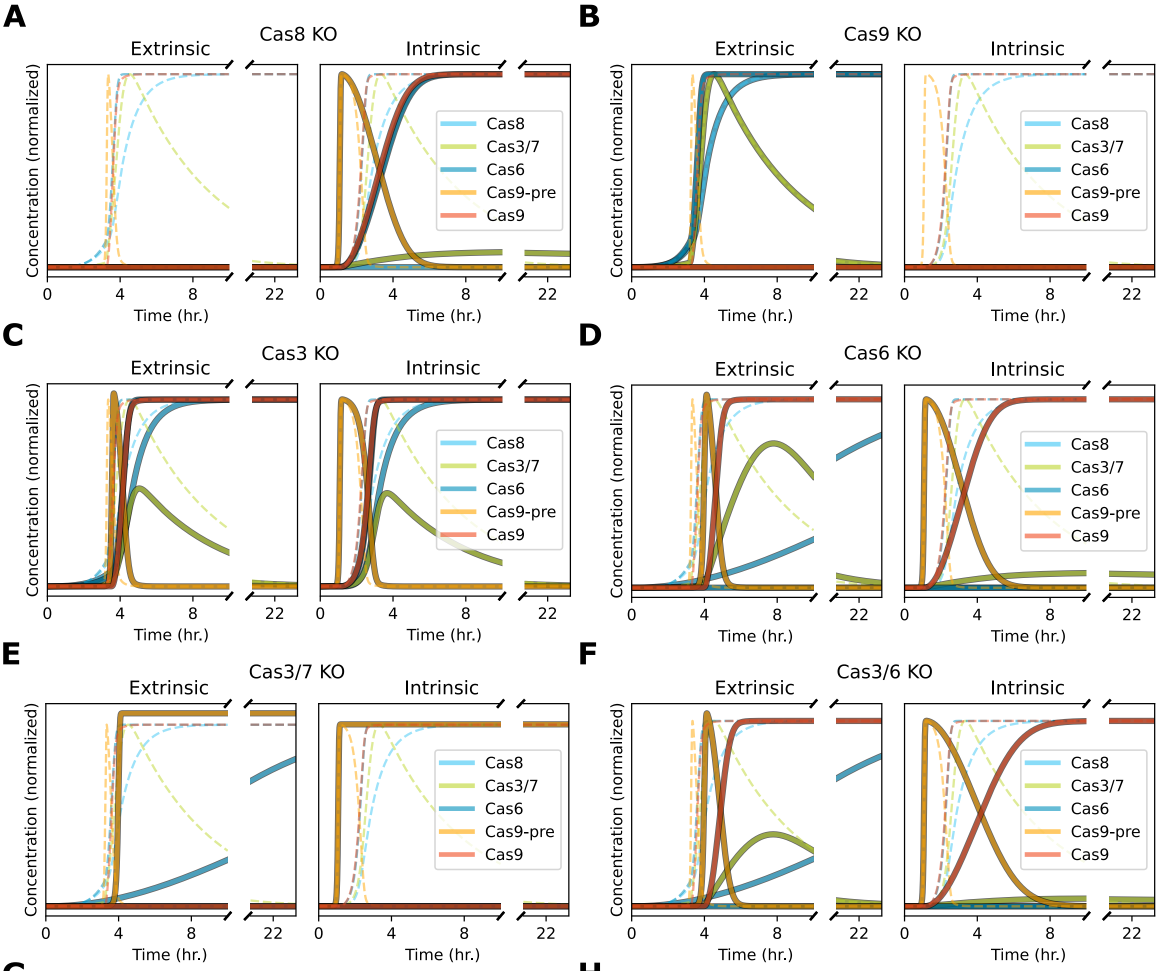


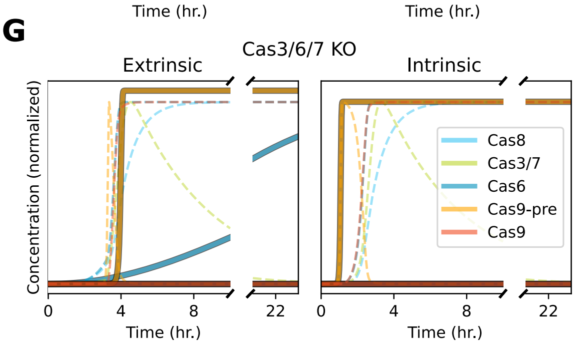


**H**

| KO | stimuli | Cas3 | Cas6 | Cas8 | Cas9-pre | Cas9-post |
|---|---|---|---|---|---|---|
| Cas8 | extrinsic | ✗ | ✗ | ✗ | ✗ | ✗ |
| | intrinsic | ✓ | ✓ | ✗ | ✓ | ✓ |
| Cas9 | extrinsic | ✓ | ✓ | ✓ | ✗ | ✗ |
| | intrinsic | ✗ | ✗ | ✗ | ✗ | ✗ |
| Cas3 | extrinsic | ✓ | ✓ | ✓ | ✓ | ✓ |
| | intrinsic | ✓ | ✓ | ✓ | ✓ | ✓ |
| Cas6 | extrinsic | ✓ | ✗ | ✓ | ✓ | ✓ |
| | intrinsic | ✓ | ✗ | ✗ | ✓ | ✓ |
| Cas3/7 | extrinsic | ✗ | ✗ | ✓ | ✓ | ✗ |
| | intrinsic | ✗ | ✗ | ✗ | ✓ | ✗ |
| Cas3/6 | extrinsic | ✓ | ✗ | ✓ | ✓ | ✓ |
| | intrinsic | ✓ | ✗ | ✗ | ✓ | ✓ |
| Cas3/6/7 | extrinsic | ✗ | ✗ | ✓ | ✓ | ✗ |
| | intrinsic | ✗ | ✗ | ✗ | ✓ | ✗ |

**Supplementary Figure 6: Simulating knock-out experiments of caspases shows agreement between predictions and western blot experiments previously performed [1, 2].** (Continued in following page)

**Supplementary Figure 6: A-G**. Time lapses of all caspases corresponding to extrinsically (left) and intrinsically (right) stimulated cells that have been simulated with different caspases knocked-out. Knock-out simulations include extrinsic caspase-8 (**A**), intrinsic caspase-9 (**B**), effector caspase-3, which is simulated as halving pro-caspase-3 initial concentration as it is redundant with effector caspase-7 (**C**), caspase-6 (**D**), both effector caspases -3 and -7 (**E**), effector caspase-3 and caspase-6 (**F**) and both effector caspases -3 and -7 as well as caspase-6 (**G**). **H**. Table summarizing the interpretation of all knock-out simulations. Each tick corresponds to a caspase that was found to be active during the simulation while crosses show caspases that were never cleaved. For example, knocking-out extrinsic (intrinsic) caspase and using extrinsic (intrinsic) stimuli shows no activation whatsoever of any caspase (**A, B, H**). Contrarily, caspase-3 KO cells show activity of every caspase as its activity is redundant with caspase-7 (**C, H**). Caspase-6 KO with and without caspase-3 simultaneous KO cells are similar and show activity of every other caspase when extrinsically stimulated while extrinsic caspase-8 can not be activated when cells are intrinsically activated (**D, F, H**). As caspase-6 is only activated by effector caspases-3 and -7, KO cells for all effector caspases with or without caspase-6 are identical. In these cases, intrinsically stimulated cells only perform a first activation of intrinsic caspase-9, which cannot fully cleave its substrates, while extrinsically stimulated ones also activate extrinsic caspase-8 (**E, G, H**).

### Supplementary Model

#### Monomers

| Monomer | Binding Sites | States |
|---|---|---|
| L | bf | |
| R | bf | |
| DISC | bf | |
| flip | bf | |
| C8 | bf | pro,A |
| BAR | bf | |
| Bid | bf | U,T,M |
| Bax | bf,s1,s2 | C,M,A |
| Bcl2 | bf | |
| CytoC | bf | M,C,A |
| Smac | bf | M,C,A |
| Apaf | bf | I,A |
| Apop | bf | |
| C3 | bf | pro,A,ub |
| C6 | bf | pro,A |
| C9 | bf | |
| PARP | bf | U,C |
| XIAP | bf | |
| Mito | bf | I,A |
| Bcl2c | b | |
| BFP | sl,bf | |
| Cit | sl,bf | |
| mKate | sl,bf | |

**Initial Concentrations**

| Species | Initial Concentration |
|---|---|
| L | 1000 |
| R | 200 |
| flip | 100 |
| C8 | 20000 |
| BAR | 1000 |
| Apaf | 1000 |
| C3 | 10000 |
| C6 | 10000 |
| PARP | 1000000 |
| XIAP | 10000 |
| Bid | 40000 |
| Bax | 100000 |
| Bcl2 | 20000 |
| Mito | 500000 |
| Smac | 100000 |
| CytoC | 100000 |
| Bcl2c | 20000 |
| dBFP | 750000 |
| dCit | 750000 |
| dmKate | 750000 |

**Reactions**

| Reaction | Forward Rate | Backward Rate |
|---|---|---|
| L + R ↔ LR | 4e-07 | 0.001 |
| LR → DISC | 1e-05 | |
| DISC + C8 ↔ DISCC8 | 1e-06 | 0.001 |
| DISCC8 → DISC + C8 | 1 | |
| C8 + Bid ↔ C8Bid | 1e-07 | 0.001 |
| C8Bid → C8 + Bid | 1 | |
| DISC + flip ↔ DISCflip | 1e-06 | 0.001 |
| BAR + C8 ↔ BARC8 | 1e-06 | 0.001 |
| Smac ↔ Smac | 0.01 | 0.01 |
| CytoC ↔ CytoC | 0.01 | 0.01 |
| CytoC + Apaf ↔ CytoCApaf | 5e-07 | 0.001 |
| CytoCApaf → CytoC + Apaf | 1 | |
| Apaf + C3 ↔ ApafC3 | 5e-09 | 0.001 |
| ApafC3 → Apaf + C3 | 1 | |
| C3 + Apaf ↔ C3Apaf | 1.3e-06 | 0.001 |
| C3Apaf → C3 + Apop | 1 | |
| Apop + C3 ↔ ApopC3 | 5e-09 | 0.001 |
| ApopC3 → Apop + C3 | 1 | |
| Apaf + XIAP ↔ ApafXIAP | 2e-06 | 0.001 |
| Smac + XIAP ↔ SmacXIAP | 7e-06 | 0.001 |
| C8 + C3 ↔ C8C3 | 1e-07 | 0.001 |
| C8C3 → C8 + C3 | 1 | |
| XIAP + C3 ↔ XIAPC3 | 2e-06 | 0.001 |
| XIAPC3 → XIAP + C3 | 0.1 | |
| C3 + PARP ↔ C3PARP | 1e-06 | 0.01 |
| C3PARP → C3 + PARP | 1 | |
| C3 + C6 ↔ C3C6 | 1e-06 | 0.001 |
| C3C6 → C3 + C6 | 1 | |
| C6 + C8 ↔ C6C8 | 3e-08 | 0.001 |
| C6C8 → C6 + C8 | 1 | |
| Bid + Bax ↔ BidBax | 1e-07 | 0.001 |
| BidBax → Bid + Bax | 1 | |
| Bax ↔ Bax | 0.01 | 0.01 |
| Bax + Bax ↔ BaxBax | 1.4e-05 | 0.001 |
| BaxBax + BaxBax ↔ BaxBaxBaxBax | 2.9e-05 | 0.0005 |
| Bax + Bcl2 ↔ BaxBcl2 | 1.4e-05 | 0.001 |
| BaxBax + Bcl2 ↔ BaxBaxBcl2 | 1.4e-05 | 0.001 |
| MatchOnceBaxBaxBax + Bcl2 ↔ MatchOnceBaxBaxBaxBcl2 | 1.4e-05 | 0.001 |
| MatchOnceBaxBaxBax + Mito ↔ MatchOnceBaxBaxBaxMito | 1.4e-05 | 0.001 |
| MatchOnceBaxBaxBaxMito → Mito | 1 | |
| Mito + Smac ↔ MitoSmac | 2.9e-05 | 0.001 |
| MitoSmac → Mito + Smac | 10 | |
| Mito + CytoC ↔ MitoCytoC | 2.9e-05 | 0.001 |
| MitoCytoC → Mito + CytoC | 10 | |
| Bid + Bcl2c ↔ BidBcl2c | 1e-06 | 0.001 |

| Reaction | Forward Rate | Backward Rate |
|---|---|---|
| BFPBFP + C3 $\leftrightarrow$ BFPBFPC3 | 2.8e-07 | 0.01 |
| C3BFPBFP $\rightarrow$ BFP + BFP + C3 | 1 | |
| CitCit + Apop $\leftrightarrow$ CitCitApop | 2.8e-07 | 0.001 |
| ApopCitCit $\rightarrow$ Cit + Cit + Apop | 1 | |
| mKatemKate + C8 $\leftrightarrow$ mKatemKateC8 | 5.4e-08 | 0.001 |
| C8mKatemKate $\rightarrow$ mKate + mKate + C8 | 1 | |
| CitCit + Apaf $\leftrightarrow$ CitCitApaf | 2e-10 | 0.001 |
| ApafCitCit $\rightarrow$ Cit + Cit + Apaf | 1 | |
